# Widespread selection against deleterious mutations in the *Drosophila* genome

**DOI:** 10.1101/2020.02.12.946392

**Authors:** Pavel Khromov, Alexandre V. Morozov

**Affiliations:** Department of Physics & Astronomy and Center for Quantitative Biology, Rutgers, The State University of New Jersey, Piscataway, NJ 08854, USA

## Abstract

We have developed a computational approach to simultaneous genome-wide inference of key population genetics parameters: selection strengths, mutation rates rescaled by the effective population size and the fraction of viable genotypes, solely from an alignment of genomic sequences sampled from the same population. Our approach is based on a generalization of the Ewens sampling formula, used to compute steady-state probabilities of allelic counts in a neutrally evolving population, to populations subjected to selective constraints. Patterns of polymorphisms observed in alignments of genomic sequences are used as input to Approximate Bayesian Computation, which employs the generalized Ewens sampling formula to infer the distributions of population genetics parameters. After carrying out extensive validation of our approach on synthetic data, we have applied it to the evolution of the *Drosophila melanogaster* genome, where an alignment of 197 genomic sequences is available for a single ancestral-range population from Zambia, Africa. We have divided the *Drosophila* genome into 100-bp windows and assumed that sequences in each window can exist in either low- or high-fitness state. Thus, the steady-state population in our model is subject to a constant influx of deleterious mutations, which shape the observed frequencies of allelic counts in each window. Our approach, which focuses on deleterious mutations and accounts for intra-window linkage and epistasis, provides an alternative description of background selection. We find that most of the *Drosophila* genome evolves under selective constraints imposed by deleterious mutations. These constraints are not confined to known functional regions of the genome such as coding sequences and may reflect global biological processes such as the necessity to maintain chromatin structure. Furthermore, we find that inference of mutation rates in the presence of selection leads to mutation rate estimates that are several-fold higher than neutral estimates widely used in the literature. Our computational pipeline can be used in any organism for which a sample of genomic sequences from the same population is available.

## 1 Introduction

Explaining the origin of genetic variation observed in natural populations is a long-standing problem in evolutionary biology. While earlier views of genome evolution were based on neutral theory, which emphasizes the role of random genetic drift in the observed patterns of intra- and interspecies variation,^1^ recent studies have questioned the applicability of the neutral theory or its nearly neutral extension^2^ to genome evolution.

One of the strongest cases for the pervasive role of natural selection in metazoans has been made in *Drosophila melanogaster*, ^3^ where population genetics modeling is enabled and guided by the availability of hundreds of sequenced genomes, including 197 from a single population in Zambia, Africa,^4, 5^ and by the functional genomics databases such as FlyBase.^6^ In particular, genomic data from *D. melanogaster* and related species was used to argue that a large fraction of the *D. melanogaster* genome, including non-coding regions, is under widespread purifying and positive selection.^7, 8^ Genetic linkage, long recognized as a key evolutionary force,^9, 10^ is also likely to play an important role in fly evolution, either through selective sweeps: hitchhiking of sites adjacent to a beneficial (adaptive) site that rises rapidly to fixation^11–13^ or through background selection: continuous generation and removal of strongly deleterious mutations by natural selection, in the presence of recombination.^14–16^

Although the evidence that selective forces and genetic linkage play key roles in *Drosophila* evolution is compelling, the exact nature and the relative contributions of the underlying evolutionary processes are less clear.^17, 18^ In particular, selective sweeps and background selection produce qualitatively similar outcomes that may be difficult to differentiate given available genomic data.^3^ A recent study has argued that background selection alone can account for a large fraction of the observed patterns of nucleotide diversity in *Drosophila melanogaster*.^19^ The study predicted nucleotide diversity at neutral sites in the presence of selection against deleterious mutations at genetically linked sites.^15, 20^ The model required several inputs: deleterious mutation rates, a parameterized distribution of deleterious selection coefficients, and the recombination frequency between the focal neutral site and the site under selection.

Here we provide an alternative approach to modeling background selection. We model the fitness landscape explicitly by assigning each allele either to a low- or a high-fitness state. The fitness difference between the two fitness states, the overall mutation rate and the fraction of alleles in the high-fitness state are inferred from rather than input into the model. To carry out the inference, we assume that the population has reached steady state and use a generalization of the neutral Ewens sampling formula^21, 22^ to fitness landscapes with multiple distinct states.^23^ As with the neutral Ewens sampling formula, basic input data consists of counts of distinct alleles in samples of aligned genomic sequences. In the absence of selection, the population maintains a steady state characterized by mutation-drift balance. As the strength of selection (i.e., the fitness difference between the two fitness states) is increased, alleles become more and more concentrated on the upper fitness plane, with each high-fitness allele subject to both neutral and deleterious mutations. Selection against deleterious mutations affects the observed frequency spectrum and the distribution of allelic counts, allowing us to carry out the inference process. We parse the *D. melanogaster* genome into non-overlapping 100-bp windows. Each window contains up to 197 aligned sequences from the Zambian fly population, providing the allelic counts that serve as input to the computational inference pipeline. Due to the complexity of the generalized Ewens sampling formula, we have opted for Approximate Bayesian Computation (ABC), a Bayesian inference method that can be used to estimate posterior distributions of model parameters.^24–26^

The key strength of the generalized Ewens sampling approach is that it goes beyond standard statistical tests of natural selection,^27^ yielding explicit estimates of key evolutionary parameters such as mutation rate and selection strength. At the same time, in contrast to the Poisson Random Field approach,^28–31^ our methodology does not assume that each nucleotide evolves independently, and therefore is capable of treating site linkage and epistasis within each genomic window. However, as with every population genetics model, several simplifying assumptions have to be made. First, the Ewens sampling approach implies that any allele can mutate into any other allele with a single mutation rate. We investigate this issue both in this work and in Ref.^23^ and conclude that systematic errors caused by this assumption are likely to be modest. Second, unlike the approach to background selection originally described in Refs.,^15, 20^ we do not treat recombination explicitly. This issue is mitigated by employing short 100-bp genomic windows (compared e.g. to 1-100 kbp windows in Ref.^19^), which however make the analysis more computationally challenging. Third, we assume that the genomic sequences in each window are in a de-labeled steady state:^22^ although the sequences keep mutating into novel alleles, the de-labeled population statistics such as the frequency spectrum and the number of distinct alleles are stationary on average. If the steady-state assumption is correct, our inference should not be affected by past expansions and contractions in the population size, such as the significant bottleneck inferred for the Zambian *D. melanogaster* population that we use in this work.^32^ Moreover, even if the population is still expanding after the bottleneck, our approach may still be applicable but will yield evolutionary parameters based on a reduced effective population size which is in turn affected by implicit bottleneck parameters.

Using the approach described above, we find that most of the fly genome evolves under selective pressure from deleterious mutations, and that for the alleles in the high-fitness state, most mutations are deleterious. The former observation is in line with the emerging consensus view on the selective constraints imposed on the evolution of the *D. melanogaster* genome,^3^ and with a recent observation of the major role of purifying selection in establishing the observed patterns of nucleotide diversity across the fly genome.^19^ Our approach thus establishes a baseline against which other selective signatures such as selective sweeps or balancing selection are to be discerned. Our methodology presents an alternative to statistical tests for selection, including background selection under recombination, and it is reassuring that it reaches broadly similar conclusions as the previous studies. Moreover, it provides an alternative to the Poisson Random Field framework as a population genetics method capable of estimating selection strength directly from single-population DNA sequence data. Finally, our approach yields a revised estimate of *θ*, mutation rates rescaled by the effective population size, suggesting that widely used neutral estimates need to be reconsidered, as neutral estimates of *θ* are systematically biased towards lower values.

## 2 Materials and Methods

### 2.1 Overview of Approximate Bayesian Computation (ABC)

ABC is an efficient inference method that can be used to estimate posterior distributions of model parameters in cases where the likelihood function is either unavailable as a closed-form expression or computationally costly to evaluate.^24–26^ Let us suppose that we have observed a statistic **x**_**0**_ and we would like to learn the distribution of model parameters ***α*** that would be consistent with the observed statistic **x**_**0**_. We also have a probabilistic model *M* that can generate the statistic of interest from the underlying parameters: ***α*** → **x**. Theoretically, it means that there exists a likelihood function *p*(**x**|***α***) but in practice either the closed form of this distribution is unknown or the computation of the likelihood is prohibitively expensive.

In such cases, Approximate Bayesian Computation (ABC) can be employed to infer the distribution of model parameters.^24–26^ ABC is based on the following rejection algorithm: First, pick a prior *p*(***α***) and sample from it to get an empirical distribution of model parameters (***α***_1_, …, ***α***_*m*_). Using the model *M*, generate the corresponding sample for the statistic of interest: (**x**_1_, …, **x**_*m*_) (i.e., generate (**x**_1_, …, **x**_*m*_) by sampling once from each *p*(**x**|***α***_*i*_), *i* = 1 … *m*). Second, choose an appropriate measure of distance *d* in the space of statistics. Then the rejection algorithm amounts to keeping only the parameters ***α***_*i*_ that generated the statistics **x**_*i*_ which satisfy *d*(**x**_*i*_, **x**_**0**_) *< ϵ* for a certain tolerance *ϵ*. The tolerance hyperparameter *ϵ* can be chosen by requiring that we keep only a certain percentage or a certain number of *m* datapoints that best match the observation.^24^ The final distributions of the model parameters retained after the rejection step constitute empirical approximations to posterior probabilities.

### 2.2 *D. melanogaster* population genomics data

In this work, we employ sequenced and aligned haploid embryo genomes from a single ancestral range population of *D. melanogaster* from Zambia, Africa, which were made available in phase 3 of the Drosophila Population Genomics Project (DPGP3).^4, 5^ Specifically, our data consists of *n* = 197 aligned sequences for chromosomes 2L, 2R, 3L, and 3R, and *n* = 195 aligned sequences for chromosome X. To summarize the observed allelic diversity in this dataset, we have divided each chromosome into 100 bp non-overlapping windows. The 100-bp window size was chosen to be able to observe non-trivial patterns of allelic diversity while minimizing the effects of recombination. We remove all windows in which more than 20% of the sequences in the alignment have at least one undetermined nucleotide (labeled as ‘N’). If the fraction of such sequences is *<* 20%, we keep the window but remove the affected sequences. We also remove all monomorphic windows as their frequency spectrum is uninformative for our analysis. This preprocessing step results in retaining 68% of all windows in chromosome 2L, 73% in 2R, 70% in 3L, 75% in 3R, and 63% in X.

For each remaining window, we compute a vector of allelic counts **n** = {*n*_1_, …, *n*_*k*_}, where *k* is the total number of groups of identical sequences in the window and *n*_*j*_ is the number of identical sequences in group *j* = (1 … *k*). Since 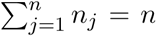, **n** can be viewed as an integer partition of *n*. As an example of the calculation of allelic counts, consider a genomic window in which out of *n* = 197 aligned sequences, 187 occur once, 3 sequences occur twice, and 1 sequence occurs 4 times. The corresponding vector of allelic counts is given by **n** = {4, 2, 2, 2, 1, …, 1}, where 1 is repeated 187 times, so that *k* = 191. Note that the same information can be conveyed with the frequency spectrum widely used in population genetics:^21, 22, 33^ **a** = (*a*_1_, …, *a*_*n*_), where *a*_*j*_ is the number of groups of identical elements of size *j*. The *a*_*j*_ counts are subject to normalization 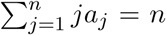. In the above example, *a*_1_ = 187, *a*_2_ = 3, *a*_4_ = 1, and the rest of *a*_*j*_’s are equal to 0. We shall use the **n** representation in the rest of the paper.

### 2.3 Application of ABC in population genetics

The main objective of this study is to infer genome-wide distributions of key evolutionary parameters such as selection strengths and mutation rates directly from sequence alignments of *D. melanogaster* genomes sampled from a single population. To this end, we use the generalized Ewens sampling formula which describes allelic diversity of a steady-state population evolving on an arbitrary fitness landscape.^23^ This is an infinite-allele model where any allele can mutate into any other allele with a single mutation rate. Unlike approaches that treat deleterious background selection with recombination,^15, 19, 20^ our method does not take recombination into account explicitly; however, in contrast to the Poisson Random Field approach,^28–31^ our methodology does not assume site independence and therefore is capable of treating linkage within each genomic window.

For a general fitness landscape with *M* distinct fitness states, the probability of the vector of allelic counts **n** is given by^23^

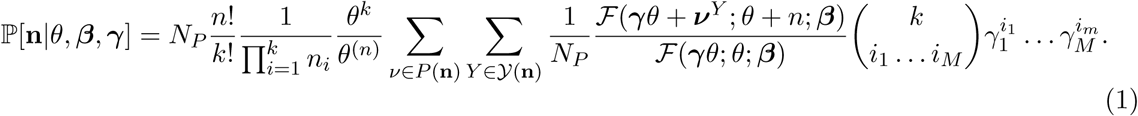

Here *θ* = *aNµ* is the rescaled mutation rate, where *N* is the effective population size, *µ* is the mutation rate per locus, and the prefactor *a* = 2 for the Wright-Fisher model^34^ and 1 for the Moran model.^35^ Fitness landscapes are encoded via a vector of selection strengths ***s*** rescaled by the population size, ***β*** = *N* ***s***, and a vector ***γ*** which determines landscape geometry by specifying the fraction of alleles (genotypes) in each fitness state. For example, with only two fitness states (*M* = 2), we assign fitness 1 to the fraction 1 − *γ* of all alleles and fitness 1 + *s* to the remaining fraction *γ*, resulting in ***β*** = (0, *Ns*) and ***γ*** = (1 − *γ, γ*). In this case, 1 − *γ* can be interpreted as a fraction of deleterious mutations for an allele in a high-fitness state. Finally, *N*_*P*_ is the total number of distinct permutations of the allelic counts, and ℱ(**a**; *b*; **z**) is the generalized confluent hypergeometric function.^23^ The double sum takes into account all the ways in which an allelic partition **n** can be distributed among *M* fitness states.

In the absence of selection, Eq. (1) reduces to the neutral Ewens sampling formula^21^ written in terms of the allelic counts *n*_*i*_:

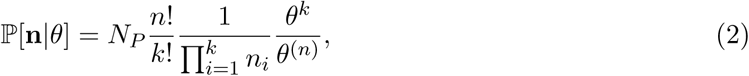

where *θ*^(*n*)^ = *θ*(*θ* + 1) … (*θ* + *n* − 1) is the rising factorial.

Additional details, along with representative examples, on how Eq. (1) is derived and evaluated, including how the double summation is performed and the numerical treatment of generalized confluent hypergeometric functions, can be found in Ref.^23^ Here it suffices to treat Eq. (1) as a “black box” function which is used in the ABC inference pipeline. Our overall objective is to learn genome-wide distributions of model parameters ***α*** = (*θ*, ***β, γ***) that appear in the generalized Ewens formula (Eq. (1)). In this case, a natural choice of the summary statistic **x** would be allelic counts **n** observed in each 100-bp genomic window. However, with *n* = 197 the total number of possible partitions is > 3 × 10^12^, too large to perform the double summation in Eq. (1). Therefore, in each window we subsample aligned sequences *B* = 10^4^ times with replacement, creating sequence samples of size *n′* = 5, for which the probabilities of the corresponding allelic counts **n***′* are amenable to evaluation using Eq. (1) or Eq. (2). In this way, we arrive at the empirical distribution ℙ[**n***′*], where 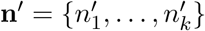 encodes all possible partitions for the smaller integer 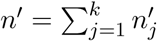. Note that our observed summary statistic **x**_**0**_ is now ℙ[**n***′*], an empirical histogram of the frequencies of allelic counts generated by subsampling alignments of *n′* = 5 sequences in each window, as described above. Since we do not have an explicit likelihood formula for this statistic, likelihood-based methods cannot be applied, whereas the ABC framework is still capable of yielding posterior distributions of model parameters.

Next, we choose model parameter priors for ABC sampling. In the case of the fitness landscape with *M* = 2 fitness states, the model parameters are ***α*** = (*θ, Ns, γ*), where *γ* is the fraction of nodes with higher fitness. We impose the following priors: log_10_ *θ* ∼ Normal(*µ* = 0, *σ* = 1), *Ns* ∼ HalfNormal(*σ* = 20) (note that if *X* is distributed according to a normal distribution with zero mean, |*X*| is distributed according to a half-normal distribution), and − log_10_ *γ* ∼ HalfNormal(*σ* = 6). These distributions reflect our prior expectations of the relevant parameter ranges. We create *m* = 10^6^ simulated empirical histograms **x**_*i*_ = ℙ[**n***′*|***α***_*i*_] (*i* = 1 … *m*) by sampling parameters ***α*** from their priors and calculating ℙ[**n***′*|***α***] via Eq. (1). We rank this dataset against the **x**_**0**_ = ℙ[**n***′*] empirical histogram of partition frequencies observed in each window, on the basis of the chi-squared statistic used as a measure of the distance between the two histograms: *d*^2^(**x**_*i*_, **x**_**0**_) = Σ_*p*_(*x*_*i,p*_ − *x*_0,*p*_)^2^*/x*_*i,p*_, where *p* labels allelic count partitions and *x*_*i,p*_, *x*_0,*p*_ are the predicted and observed frequencies of the allelic partition *p*. We set ABC tolerance *ϵ* by ranking all 10^6^ simulated histograms against the histogram observed in a given window and keeping 10^2^ histograms with the smallest *d*^2^ score. We typically use median values to summarize the resulting posterior distributions of the model parameters, since they are less sensitive to outliers.

### 2.4 Recombination simulations

Population dynamics under mutation, selection and recombination is modeled using the Moran process.^35^ The population consists of *N* = 10^3^ sequences with alphabet *A* = 4 and length *L* = 10 sites. Each evolutionary simulation starts with a randomly generated population. Subsequent generations are obtained using the following rules:

- Two distinct parental sequences *s*_*i*_ = (*p*_1_, …, *p*_*L*_) and *s*_*j*_ = (*q*_1_, …, *q*_*L*_), *i* ≠ *j*, are selected from the population *s*_1_, …, *s*_*N*_ with weights proportional to their fitness.
- A random number *r* is uniformly sampled in the (0, 1) range. If *r* is less than the recombination rate *ρ*, an offspring sequence (*p*_1_, …, *p*_*b*−1_, *q*_*b*_, …, *q*_*L*_) is formed using the parent sequences *s*_*i*_ and *s*_*j*_, where the breakpoint *b* is uniformly sampled from 2 ≤ *b* ≤ *L* − 1. If *r* > *ρ*, recombination does not occur and the offspring is a copy of the first parent *s*_*i*_.
- The offspring undergoes a single-point mutation with probability *µ* at a randomly chosen site 1 ≤ *k* ≤ *L*.
- The population is uniformly sampled to remove a single sequence. The offspring sequence is then added to the population, keeping the total population size constant.

The above steps are repeated until the steady state is reached. Once in steady state, the population is sampled 10^4^ times, skipping 1*/µ* generations between consecutive samples.

## 3 Results

### 3.1 Validation of the ABC inference pipeline using synthetic data

In order to validate our ABC approach to the genome-wide inference of evolutionary parameters, we first test our pipeline on simulated data. We use Eq. (1) with *M* = 2 fitness states to generate sampling probabilities for *n′* = 5 on a grid of rescaled selection coefficients, *Ns*, and the fraction of alleles with fitness 1 + *s, γ*, for three values of the rescaled mutation rate *θ*. We use these probabilities directly as input to the ABC inference pipeline, which returns predictions for these 3 parameters. Note that this procedure is equivalent to producing an infinitely large sample of sets of 5 aligned sequences, so that we are not testing how sampling noise affects the accuracy of predictions (our sample size in each *D. melanogaster* genomic window, *B* = 10^4^, is sufficiently large to minimize the sampling noise effects). Since we know the exact values of all 3 evolutionary parameters ***α*** = (*θ, Ns, γ*), we can systematically evaluate the accuracy of ABC inference across biologically relevant parameter ranges (Fig. 1). The same set of *m* = 10^6^ simulated empirical histograms ℙ[**n***′*|***α***_*i*_] (*i* = 1 … *m*) is used here as in subsequent inference on fly genomic data (see Materials and Methods for details).

**Figure 1:**
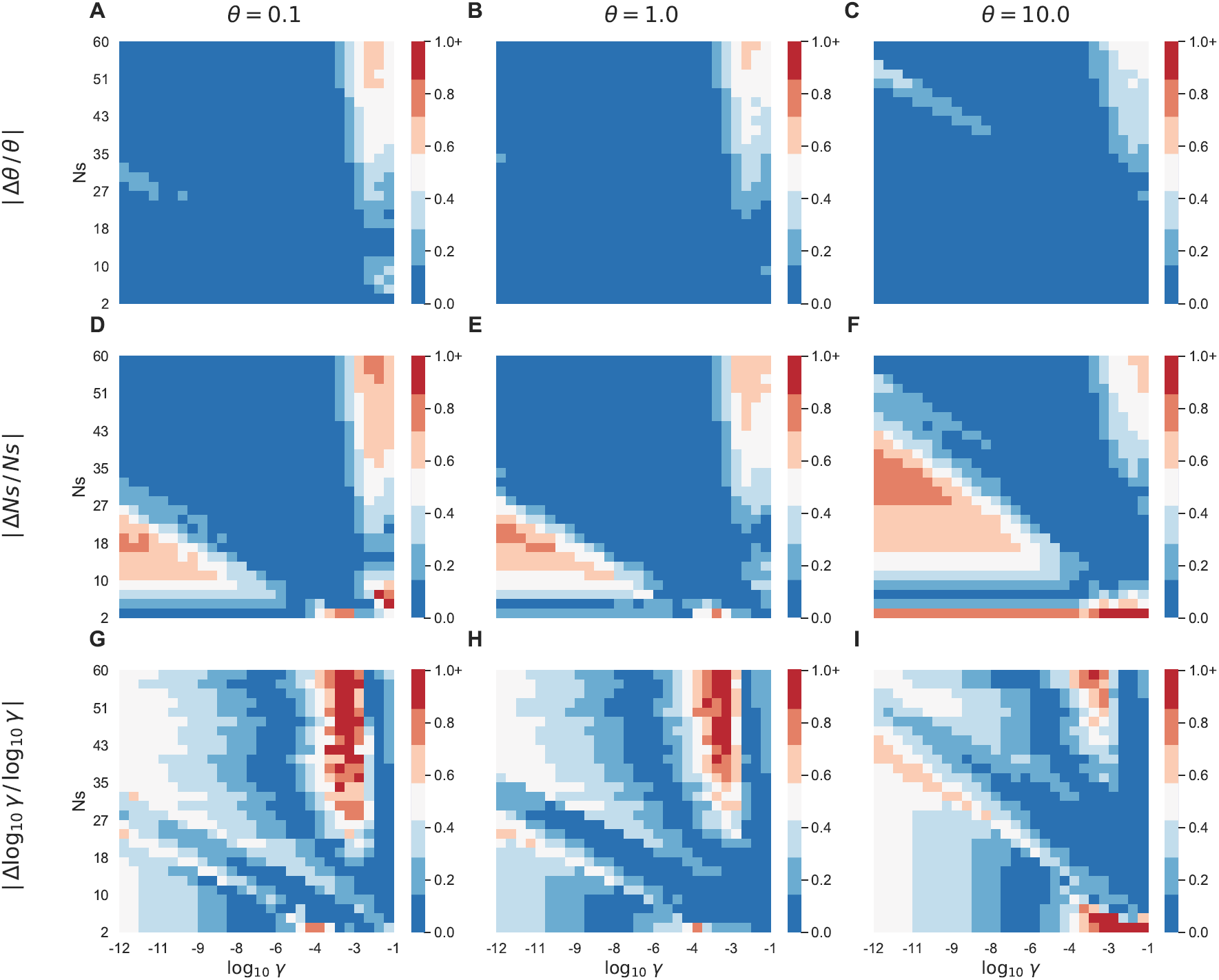
Accuracy of ABC inference with selection on synthetic data. Shown are relative errors Δ*x/x* = (*x*_inferred_ − *x*_true_)*/x*_true_ (where *x*_inferred_ is taken to be equal to the median value of the corresponding posterior distribution) for the rescaled mutation rate *θ* (A, D, G), the rescaled selection coefficient *Ns* (B, E, H), and the log-fraction of alleles with fitness 1 + *s*, log_10_ *γ* (C, F, I). All relative errors are calculated for the same ranges of *Ns* and *γ* and for *θ* = 0.1, 1.0, 10.0 corresponding to the monomorphic, intermediate, and polymorphic regimes.

Overall, the accuracy of our inference procedure is encouraging in this idealized setting, where the same model is used to generate the data and infer model parameters from it. In particular, we can reliably estimate mutation rates *θ* almost everywhere within the parameter ranges shown in Fig. 1A,D,G: the average relative error for *θ* inference is ≈ 0.09 over all 3 values of *θ*, while the maximum error is ≈ 0.66. For *Ns* and *γ*, there are parameter regions where the predictions yield significant errors, e.g. in the triangular-shaped area in Fig. 1B,E,H. It appears that the models are degenerate in these regions, such that multiple sets of parameters can fit input sampling probabilities in the *n′* = 5 histograms. As a result, the average relative error for *Ns* inference is ≈ 0.80, while the maximum error is ≈ 14.8. With log_10_ *γ* inference, the most prominent area where the ABC inference procedure yields significant errors is the vertical stripe in the large *γ*, large *Ns* quadrant (Fig. 1C,F,I). The area of the stripe shrinks as the population becomes more polymorphic. As with the *Ns* inference, the sampling probabilities within the stripe can be fit using multiple sets of parameters, so that the original parameter set is difficult to recover. The average and the maximum relative errors for log_10_ *γ* inference are ≈ 1.1 and ≈ 72.2, correspondingly.

We conclude from these numerical experiments that we should be able to infer the values of mutation rates from genomic data with a reasonably high degree of accuracy. Moreover, our predictions of selection coefficients, although subject to larger errors, should be accurate enough to serve as a test for the presence of natural selection signatures in genomic data. Finally, the predictions of the fraction of alleles with the higher fitness, *γ*, is the least reliable but also the least informative, since it depends on the assumption of two distinct fitness states. The structure of realistic fitness landscapes is likely to be considerably more complicated. Overall, the prediction accuracy is determined by which regions of the parameter space correspond to *D. melanogaster* genomic data.

To investigate the role of selection in predicting mutation rates, we have also carried out ABC inference using the neutral Ewens formula, Eq. (2), instead of its generalized version with selection, Eq. (1), to generate *m* = 10^6^ histograms ℙ[**n***′*|*θ*_*i*_] using the values of *θ*_*i*_ (*i* = 1 … *m*) previously employed for ABC inference with selection (Fig. 2). Since the input sampling probabilities are the same as in the previous test and therefore were generated under selection, we expect neutral inference to perform less well than the full treatment with selection. Indeed, considerable errors in *θ* predictions are produced under the neutral assumption in all three mutation regimes (Fig. 2A-C). The errors are small only in the regions where selection is weak and the fraction of high-fitness alleles is small: both of these factors make the mutational forces more dominant and the system is effectively neutral. We conclude that when population evolves under selection which is strong enough not to be dominated by mutational effects, neglecting selective forces (i.e., assuming neutrality) may easily lead to considerable errors in the inferred mutation rates.

**Figure 2:**
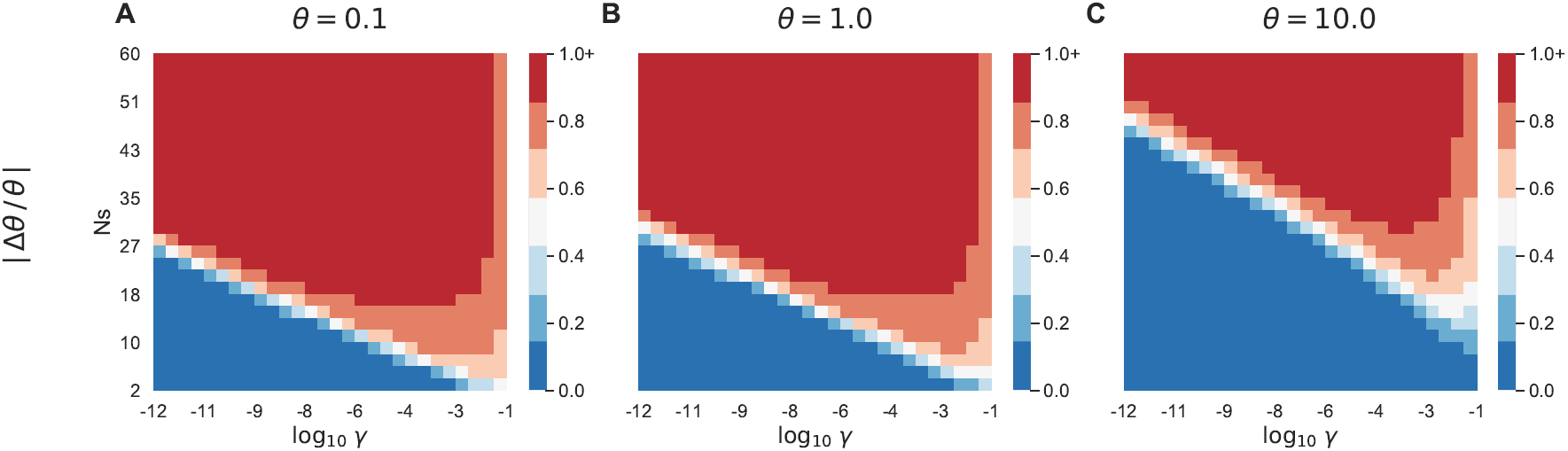
Accuracy of neutral ABC inference on synthetic data. Shown are relative errors Δ*x/x* = (*x*_inferred_ − *x*_true_)*/x*_true_ (where *x*_inferred_ is taken to be equal to the median value of the corresponding posterior distribution) for the rescaled mutation rate *θ* (A, B, C). All relative errors are calculated for the same ranges of *Ns* and *γ* as in Fig. 1 and for *θ* = 0.1, 1.0, 10.0 corresponding to the monomorphic, intermediate, and polymorphic regimes.

### 3.2 Frequencies of allelic counts and ABC inference in the presence of recombination

Evolution of genomic sequences in *D. melanogaster* is subject to homologous recombination.^36–41^ Since our ABC inference pipeline does not include recombination explicitly (instead, we rely on the small window size, *L* = 100 bp, to minimize its effects in genome-wide analysis), we thought to investigate the influence of recombination on allelic count frequencies using a simple model system. To this effect, we have simulated a population of sequences with *L* = 10 sites subject to single-point mutation, recombination, and genetic drift (see Materials and Methods for details). Thus, each site in our population is equivalent to 10 bp in genomic windows. Since *θ* per bp is less than 0.01 per bp in *D. melanogaster* according to standard infinite-sites-based estimators^41^ and therefore less than 1.0 per 100-bp genomic locus, we thought to investigate *n′* = 5 allelic count probabilities for *θ* = *Nµ* = (0.1, 1.0, 10.0) in our simulations.

Note that since the rate of spontaneous point-mutation events is ≈ 5 − 6 × 10^−9^ per nucleotide per generation according to mutation-accumulation experiments,^42, 43^ the effective population size in the Zambian population under investigation is expected to be around 10^6^ individuals,^32^ three orders of magnitude larger than the *N* = 10^3^ population size we were able to implement in our simulations. Finally, fine-scale predictions of recombination rates using two *D. melanogaster* populations, one from North America and the other from Africa, yield *ρ/µ* estimates in the ≈ 2 – 4 range.^41^ Correspondingly, we have investigated *ρ/µ* = (0, 1, 2) cases for each of the three *θ* values mentioned above and for three values of selection strength: *Ns* = (0, 6, 13) (Fig. S1). For each set of parameter values, we have compared theoretical predictions for *n′* = 5 allelic frequencies (Eq. (1)) with steady-state numerical simulations. The *ρ/µ* = 0 case with no recombination highlights the difference between the Ewens sampling approach, which assumes that each locus (i.e., nucleotide sequence in a 100-bp genomic window) can mutate into every other locus, and the numerical simulations in which only single-point mutations are allowed (see Ref.^23^ for further analysis of this assumption).

Overall, we find reasonable agreement between predicted and observed allelic count frequencies. In the monomorphic limit (*θ* = 0.1), the agreement is very good for all values of *Ns* and *ρ/µ* (Fig. S1A-C), indicating that the Ewens sampling approach would benefit from parsing the genome into even shorter, 10 − 20 bp genomic windows that are characterized by *θ* values of similar magnitude. The agreement is significantly worse in the *θ* = 1.0 regime when selection is present (Fig. S1D-F), indicating failure of the assumptions inherent in the Ewens sampling formula rather than the effects of recombination, which appear to be secondary in this case. The situation is reversed in the *θ* = 10.0 regime (Fig. S1G-I), with recombination effects becoming more prominent. At the same time, deviations between generalized Ewens formula predictions and the *ρ/µ* = 0 simulation are minimal in this regime, even in the presence of selection.

To investigate the effect of the observed discrepancies in allelic count frequencies on evolutionary parameter inference, we have used the *n′* = 5 allelic count frequencies from Fig. S1 as input to our ABC inference pipeline (Table S1). We find that *θ* is predicted very accurately for all parameter combinations, with the largest discrepancies observed when *θ* = 10.0 and *ρ/µ* = 1 or 2. For the rescaled selection coefficient *Ns*, the algorithm tends to predict non-zero values for a small fraction of alleles (e.g., *Ns* = 4.01 and *γ* = 2.67 × 10^−9^ in the case of neutral evolution with *θ* = 0.1 and *ρ/µ* = 2; Table S1, subsection A). In the ideal situation where, as in Fig. 1, Eq. (1) is used to both produce the *n′* = 5 allelic count frequencies and carry out ABC inference from them, the algorithm is able to make reasonably accurate predictions of both the selection strength and the fraction of viable alleles (cf. Theory columns in each subsection of Table S1).

Predictions are also accurate in the *θ* = 0.1 regime when allelic count frequencies from simulations with recombination are used as input to the ABC inference pipeline. However, the situation becomes more complicated when mutation rates increase: for example, in the *θ* = 1.0 regime predicted *Ns* values alone are not a reliable indicator of selection strengths unless they are supplemented by the values of *γ*, which are much higher in the *Ns* = 6 and 13 cases compared to neutral evolution (Table S1, subsections D,E,F). Even considered together, the *Ns* and *γ* predictions become unreliable in the *θ* = 10.0 regime (Table S1, subsections G,H,I), indicating that we cannot distinguish selection strength differences of 𝒪(1) in this case. In summary, our analysis indicates that it is preferable to combine evidence from both *Ns* and *γ* predictions, and that it is prudent to limit the observations to qualitative conclusions such as presence or absence of selection, especially in windows with *θ* ≳ 10.0.

### 3.3 Evolutionary parameter inference in *D. melanogaster*

#### Inference of mutation rates

We have carried out genome-wide inference of rescaled mutation rates using several alternative methods. First, we use standard mutation rate estimators due to Watterson,^44^ which is based on the number of segregating sites observed in each window, and due to Tajima,^45^ which is based on the average number of nucleotide differences between all pairs of sequences. The normalized difference of the two estimators defines Tajima’s D statistic widely used in tests for selection.^27, 46^ For each valid 100 bp window with *n* aligned unambiguous sequences (see Materials and Methods for details), Watterson estimator is given by *θ*_*W*_ = *S*_*n*_*/H*_*n*−1_, where *S*_*n*_ is the number of segregating sites and 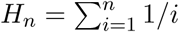 is the harmonic number. Tajima estimator, *θ*_*T*_, is simply the average number of polymorphisms (nucleotide differences) in all pairwise alignments generated by *n* aligned 100-bp sequences in each window. Tajima and Watterson estimators are compared with two ABC-based inferences of *θ*, one employing the neutral formula (Eq. (2); 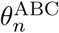) and the other one the formula with selection (Eq. (1); 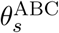) (Fig. 3, Fig. S2).

**Figure 3:**
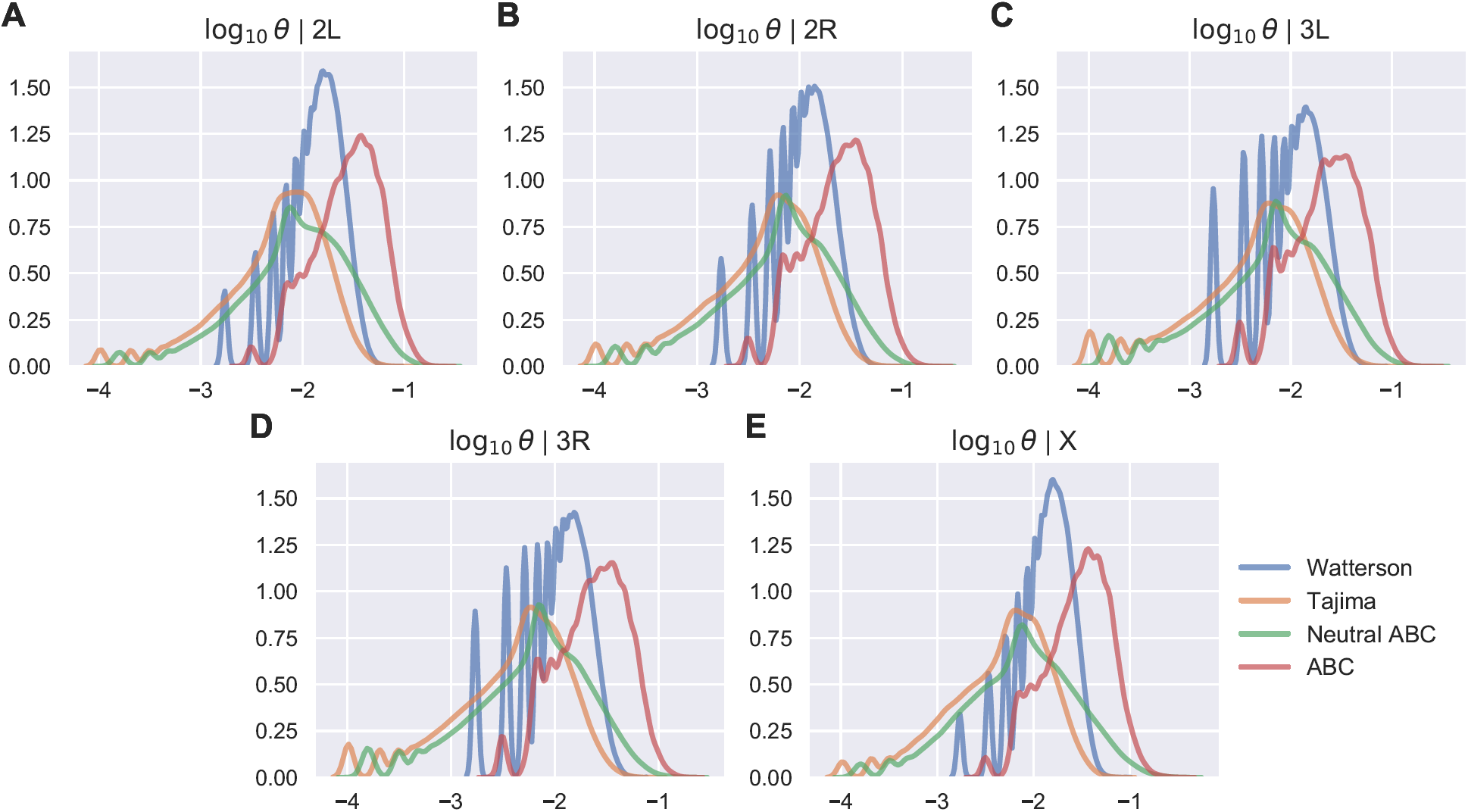
Chromosome-wide distributions of mutation rates. Shown are the histograms of Watterson (*θ*_*W*_, blue), Tajima (*θ*_*T*_, orange), ABC neutral (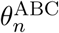, green) and ABC with selection (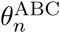, red) estimators of the logarithm of the mutation rate per bp rescaled by the population size, log_10_ *θ*, for chromosomes 2L (A), 2R (B), 3L (C), 3R (D), and X (E). The two ABC estimators are represented by their median values in each genomic window.

First of all, we observe that, surprisingly, Tajima’s estimator of the mutation rate, *θ*_*T*_, is strongly correlated with the neutral ABC estimate, 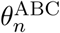. In fact, the genome-wide linear correlation between these two approaches is *r* = 0.98 (Fig. S2D). Although the two approaches are mathematically different (using pairwise polymorphisms in the former and allelic counts in the latter), they yield very similar mutation rate estimates. In contrast, the correlation coefficient between *θ*_*W*_ and *θ*_*T*_ is much smaller (*r* = 0.77; Fig. S2A) and *θ*_*T*_ − *θ*_*W*_ *<* 0 in most cases, indicating that Tajima’s D statistic, which is ∼ *θ*_*T*_ − *θ*_*W*_, is affected by selection against genotypes carrying deleterious mutant alleles (for the purposes of interpreting this test, we assume that the Zambian *D. melanogaster* population under consideration is approximately stable in size). Indeed, the distribution of Tajima’s D statistic for each chromosome is skewed towards negative values and its magnitude strongly suggests selective effects in a significant fraction of windows (Fig. S3). This can be explained by the fact that the number of segregating sites on which Watterson’s estimator is based ignores the frequency of mutations and therefore is expected to be more strongly affected by the existence of rare deleterious mutants than the average number of pairwise nucleotide differences.^45^

Both *θ*_*W*_ and 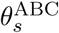 tend to predict consistently higher values of mutation rates than the neutral ABC estimate (Fig. 3). This is clearly seen when inferred mutation rates are plotted along each chromosome (Fig. 4): ABC inference with selection predicts the highest mutation rates, followed by Watterson’s estimate. However, *θ*_*W*_ is only moderately correlated with the ABC estimate under selection, 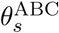 (*r* = 0.65, Fig. S2C), indicating that both estimates are affected by selective forces in somewhat different ways. This is not surprising since, as mentioned above (and unlike ABC inference with selection), Watterson’s estimator ignores the frequency of mutations. Remarkably, ABC inference with selection produces distributions of mutation rates that are nearly identical from chromosome to chromosome, indicating that the inference process is dominated by global polymorphism patterns rather than chromosome-specific features (Fig. 5A). To ensure that we do not lose information by focusing on the median values of rescaled mutation rates in each genomic window and to estimate the uncertainty of our predictions, we have constructed the genome-wide posterior probability by combining data from all windows (Fig. S4A). We find that the genome-wide distribution is essentially unimodal, with the shape similar to those seen in Fig. 5A and with predicted log_10_ *θ* values predominantly concentrated in the [−2.3, −0.8] range.

**Figure 4:**
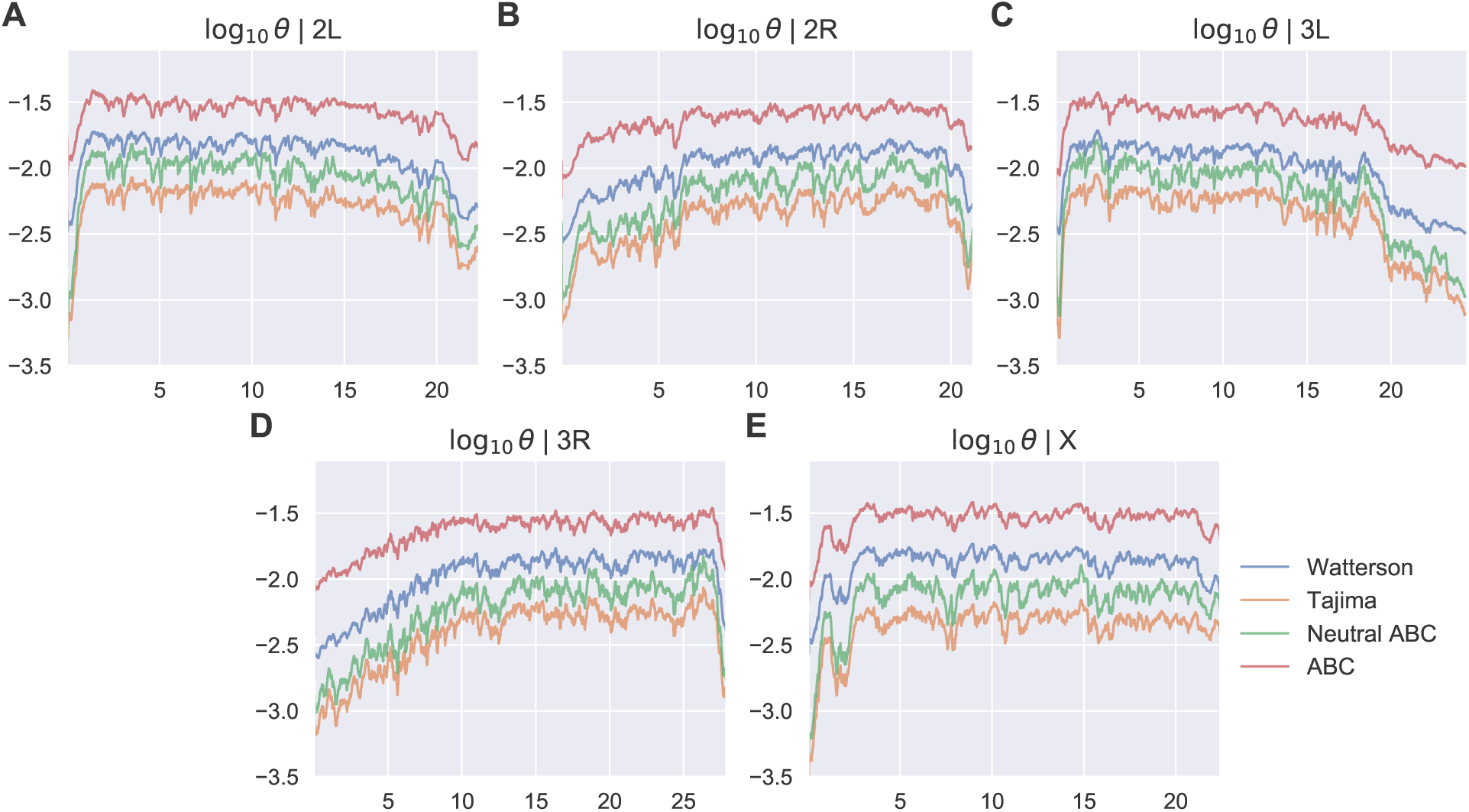
Inferred mutation rates along *D. melanogaster* chromosomes. Shown are Watterson (*θ*_*W*_, blue), Tajima (*θ*_*T*_, orange), ABC neutral (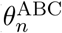, green) and ABC with selection (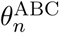, red) estimators of the logarithm of the mutation rate per bp rescaled by the population size, log_10_ *θ*, for each 100-bp window, for chromosomes 2L (A), 2R (B), 3L (C), 3R (D), and X (E), plotted vs. the genomic coordinate along each chromosome (in Mbp). The two ABC estimators are represented by their median values in each genomic window. All plotted values were smoothed with an exponentially weighted moving average with the center of mass of 1,000 windows, such that the exponential parameter *α* ≃ 10^−3^.

**Figure 5:**
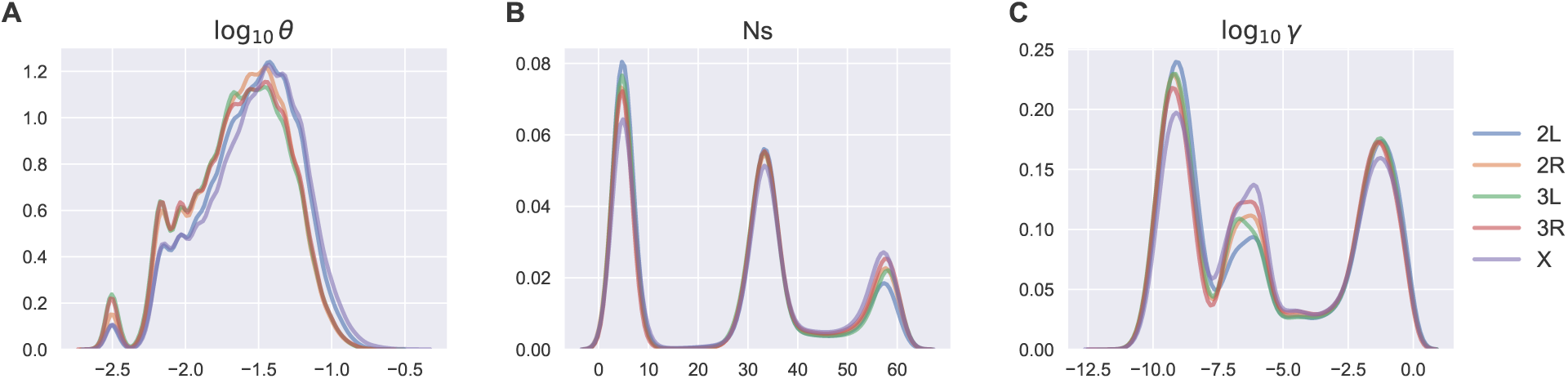
ABC inference of mutation rates, selection coefficients, and fraction of viable genotypes. Chromosome-wide distributions of log_10_ *θ* per bp (A), *Ns* (B) and log_10_ *γ* (C) based on the generalized Ewens formula with selection (Eq. (1)). All predicted quantities are represented by their median values in each window. The curves for each chromosome represent a smoothed trace of a normalized histogram.

#### Inference of selection strengths and the fraction of viable genotypes

We observe three distinct peaks of selection strengths in each chromosome: weak selection (peak 1; *Ns <* 15), intermediate selection (peak 2; 15 ≤ *Ns* ≤ 45), and strong selection (peak 3; *Ns* > 45) (Fig. 5B). As with the rescaled mutation rates, the peak structure is similar in all chromosomes. Thus ABC inference predicts that most of the fly genome evolves under selective constraints, in accordance with previous studies (see Ref.^3^ for a comprehensive review). Interestingly, three distinct peaks are also observed in the genome-wide posterior probability of selection strengths (Fig. S4B); the probability that *Ns* > 1 is 94.2% according to this distribution. The fraction of genotypes in the high-fitness state (which we shall refer to as viable genotypes), *γ*, or, alternatively, the fraction of neutral mutations for a viable allele, also exhibits a characteristic peak structure which is fairly similar for all chromosomes (Fig. 5C). However, this structure is not observed in the genome-wide posterior distribution, which is bimodal with a narrow peak in the [-3.0,0.0] range and a much broader peak in the [-20.0,-3.0) range (Fig. S4C).

To investigate whether sequences in different fitness peaks correspond to distinct distributions of mutation rates and fractions of viable genotypes, we have divided all windows into 3 classes according to selection strength (Fig. S5). We observe that mutation rates do not correlate strongly with *Ns* peak identity, although sequences with intermediate selection strengths do tend to have somewhat higher mutation rates (Fig. S5A). In contrast, fractions of viable genotypes are partitioned by selection strength, with the sequences under strong selection characterized by intermediate values of log_10_ *γ* (Fig. S5C). In the light of our previous discussion of prediction accuracy on synthetic data with and without recombination (Fig. S1, Table S1), the intermediate-selection peak may be the most reliable since it is accompanied by sizable values of log_10_ *γ*. With peaks 1 and 3, we cannot rule out the possibility that in some genomic windows, similar to predictions in Table S1, neutral evolution is in fact modeled by non-zero selection coefficients accompanied by low values of *γ*. In addition, selection strengths in peak 1 may be insufficient to reliably rule out the no-selection scenario. Finally, we note that plots of *Ns* and log_10_ *γ* vs. chromosome coordinates show no easily identifiable trends, except for the higher values of *Ns* accompanied by somewhat lower values of log_10_ *γ* in both sub-telomeric regions (Figs. S6,S7).

Next, we have investigated the nature of genomic sequences that belong to the three selection peaks. We find that, as might be expected, the strength of selection is inversely correlated with the number of mutations observed in corresponding genomic sequences. Indeed, sequences evolving under the strongest selection (peak 3) are significantly less polymorphic than sequences predicted to be under weak selection (peak 1), with sequences in peak 2 occupying an intermediate position (Fig. 6A). Besides the number of mutations per sequence, we have considered the distribution of allelic partitions in *n′* = 5 sequence alignments used in our inference procedure (Materials and Methods) (Fig. 6B). We observe that *n′* = 5 allelic counts that correspond to sequences under the strongest selective constraint (peak 3) are generated by sequences that are either all identical in the alignment ({5}) or with a single different sequence ({4, 1}). In contrast, sequence alignments in peak 1 are predominantly polymorphic, and sequence alignments in peak 2 occupy an intermediate position. Since mutation rates are predicted to be polymorphic for most windows (i.e., log_10_ *θ* + 2 > 0, where *θ* is the rescaled mutation rate per bp) regardless of their peak identity (Fig. S5A), the number of mutations in the alignment is, according to the ABC inference pipeline, indicative of selective constraints rather than the monomorphic limit.

**Figure 6:**
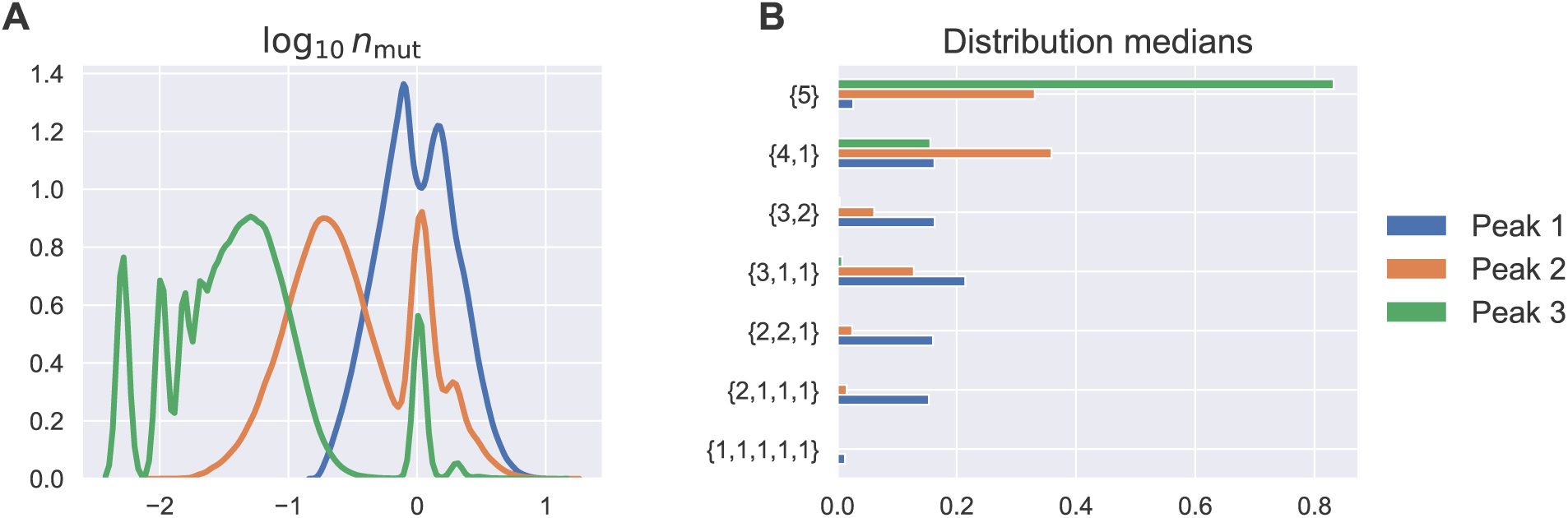
Polymorphisms in sequence alignments and selection strength. Histograms of the average number of mutations per sequence, *n*_mut_, for 100-bp genomic windows associated with different selection peaks (Fig. 5B, Fig. S5B) (A). Median values of the distribution of *n′* = 5 allelic counts for all windows in the three selection peaks from Fig. 5B (B). In both panels, data from all chromosomes is combined. In panel B, each 100-bp window has a set of 10^2^ ℙ[**n***′*|***α***_*i*_] distributions corresponding to 10^2^ sets of model parameters ***α***_*i*_ with the smallest *d*^2^ score (see Materials and Methods for details). These sets of histograms are combined into a single dataset for all windows that belong to a given selection peak and median values of the frequency distribution for each allelic configuration in the *n′* = 5 partition are reported.

It would be natural to expect that sequences under stronger selective constraints are predominantly associated with functional genomic regions, such as coding sequences and promoters. However, we do not find any correlation between the average number of mutations per sequence in each 100-bp window and its location within either a genic or an intergenic region (Fig. S8; we employ FlyBase annotation v. 6.29 to map functional regions^6^). In fact, the distributions of the average number of mutations in genes and intergenic regions are remarkably similar for each chromosome (qualitatively similar results are obtained when considering exons and introns separately; data not shown). Moreover, distributions of the average number of mutations sorted by selection peaks in Fig. S5B are strongly overlapping in each chromosome (Table S2). We conclude that sequences under weak, intermediate and strong selection are distributed throughout the fly genome in a way that is independent of their standard functional annotation.

Finally, since our ABC inference pipeline produces sizable errors in some areas of the (*θ, Ns, γ*) parameter space (cf. Fig. 1), it is possible that our results are affected by inaccuracies in ABC computational predictions. However, a comparison of the ranges of predicted parameters in Fig. S5 with the prediction errors on synthetic data in Fig. 1 shows that most of our predictions are not concentrated in the problematic regions of the parameter space. For example, windows in peak 3 have 0.3 ≲ *θ* ≲ 10, 45 ≲ *Ns* ≲ 60, and −8 ≲ log_10_ *γ* ≲ −5. A comparison with the error plots in Fig. 1 shows that we can expect excellent accuracy for *θ* and *Ns* inference and reasonable accuracy for log_10_ *γ* inference within these ranges. The same is true, by and large, of the other two peaks. We conclude that the ABC inference procedure applied to *D. melanogaster* genomic data has sufficient internal consistency for at least qualitative conclusions regarding the magnitude of mutation and selection forces. This observation does not however preclude the possibility that our results are affected by the phenomena that are not explicitly included into the ABC model formulated above, such as recombination,^36–41^ demographic effects,^32, 47^ and the assumption that any sequence in the 100-bp window can mutate into every other sequence.^21^

## 4 Discussion and Conclusion

In this work, we have developed a novel computational approach to simultaneous genome-wide inference of mutation rates, selection strengths and the average fractions of beneficial, deleterious and neutral mutations per allele. The approach is based on applying Approximate Bayesian Computation^24–26^ to the Ewens sampling formula which we have previously generalized to evolution under selection.^23^ The generalized Ewens sampling formula provides an explicit closed-form solution for the probability of each partition of *n* alleles (for example, aligned sequences in a genomic window) into allelic counts. However, it is cumbersome to implement, requiring (i) a partition of *n* aligned sequences into all possible allelic partitions and (ii) for each allelic partition, a sum over all the ways in which the partition can be distributed among different fitness states. The ABC inference pipeline alleviates these computational difficulties since it was specifically designed for cases where the probability of the statistic of interest is either not a closed-form expression or computationally costly to evaluate.

Furthermore, we have assumed that all alleles can adopt either a low- or high-fitness state, so that, with a sufficient fitness difference between the two states, sequences in the population will predominantly concentrate in the high-fitness state and, for such sequences, the newly arising mutations will be either neutral or deleterious. In this aspect, our fitness landscape conforms to a central tenet of the neutral theory which assumes that the contribution of beneficial mutations can be ignored.^1^ Note, however, that the magnitude of the difference between the two fitness states is inferred from the data rather than imposed, enabling us to differentiate between the strictly neutral scenario and its generalization to both deleterious and neutral evolutionary dynamics. Finally, the Ewens sampling approach is based on the steady-state assumption: although specific sequences that make up the evolving population change, the “de-labeled” statistics such as the average number of distinct alleles in the population is time-independent.^21, 33^ Note that a generalization to more than two fitness states is not likely to be qualitatively different since the population will always adopt the highest-fitness configuration in steady state, with the mutational load due to deleterious mutations into all lower-fitness fitness.

We have applied the ABC inference approach to study selective constraints on the genomic evolution of *D. melanogaster. D. melanogaster* is a key model organism in modern genetics and as a result evolution of fruit fly populations in the wild has received considerable attention in the population genetics community, both experimentally and computationally. In particular, a considerable number of fly genomes have been sequenced, aligned and functionally annotated to a common standard in a large-scale effort.^4, 5^ Specifically, phase 3 of the *Drosophila* Population Genomics Project (DPGP3) has provided 197 haploid embryo genomes from a single *D. melanogaster* population in Zambia, Sub-Saharan Africa. *D. melanogaster* likely originated in the Sub-Saharan region,^48^ so that the genome sample is from the species’s ancestral range. This data provides a rich collection of polymorphisms and allelic counts in a single fruit fly population. The allelic counts serve as input to the ABC inference pipeline developed in this work. To carry out ABC analysis, we have parsed the *D. melanogaster* genome into 100-bp non-overlapping windows. The size of the windows was chosen to minimize the effects of recombination, which is not explicitly treated in the Ewens sampling framework, while still dealing, in each window, with a polymorphic sample that provides informative allelic counts.

Furthermore, linkage between mutations that belong to the same window is fully taken into account, going beyond the other major assumption of the neutral theory, that positive and negative selection at linked loci does not affect the dynamics of neutral alleles.^1^ As a result, our approach is closer to the background selection framework, which explicitly treats the effects of recombination and linkage but puts emphasis on negative rather than positive selection.^14–16, 19^ Similar to background selection, our computational procedure can be viewed as a baseline model, deviations from which would be indicative of positive selection events such as selective sweeps. We find that, consistent with previous studies (reviewed in Ref.^3^), a large fraction of the *Drosophila* genome appears to evolve under selective constraints. Similar to previous work,^19^ we find that purifying selection can explain the observed patterns of nucleotide diversity in the *Drosophila* population under consideration. The major role of deleterious mutations is expected given that deleterious and neutral mutations are typically much more numerous than beneficial ones.^49^ We observe that sequences under selective constraints are not preferentially associated with coding regions or other functional elements, or with centromeric or telomeric positions (although selection does appear to be stronger at sub-telomeric regions, Fig. S6), and are instead distributed evenly throughout the genome. All sequences under selection are grouped into three distinct peaks, with weak, intermediate, and strong selection (Fig. 5B). The peaks of selection strength correlate with the total number of polymorphisms observed in a genomic window and with the frequencies of allelic counts, with sequences under weaker selection generally being more polymorphic (Fig. 6). These global constraints may reflect the need to maintain nucleosome positioning^50^ or higher-order chromatin structure,^51^ or other universal constraints whose exact nature is currently unclear.

The ability to treat linkage and epistasis within genomic windows of arbitrary width (constrained only by computational considerations) provides a substantial advantage over the Poisson Random Field approach,^28–31^ which can infer the strength of selection (rather than merely detect its presence) but is unable to account for linkage between sites. In addition to providing a quantitative Bayesian estimate of selection strength, our inference pipeline yields simultaneous estimates of population-size-rescaled mutation rates and of the fractions of neutral, deleterious and beneficial mutations for each high- and low-fitness allele in a steady-state population. Overall, our *θ* estimates yield 2-3 fold higher values genome-wide compared to standard neutral estimates^44, 45^ and our own ABC estimate without selection (Fig. 4). We conclude that ignoring selection against deleterious mutations leads to consistent underestimation of effective population sizes. Finally, we find that as a rule, there are many more deleterious than neutral mutations available to an allele in a high-fitness state. Interestingly, it is the alleles subjected to intermediate levels of selection that are the most robust to mutations (i.e., have the largest number of neutral mutations available to them) (Fig. S5C).

Our inference relies on the steady-state assumption and therefore our estimates may become inaccurate if the population is in the process of expansion or contraction. However, our framework should be able to account for past changes in the population size, such as bottlenecks, through adjusting the effective population size. *Drosophila* demographics is a potential compounding factor because not only derived fruit fly populations have been associated with severe bottlenecks,^47^ but the ancestral range population in Sub-Saharan Africa is also predicted to have undergone a significant bottleneck.^32^ These past events should reduce the magnitude of the effective population size in our framework; our predictions of *θ* (and using mutation rates per nucleotide from previous mutation-accumulation studies^42, 43^) yield *N*_eff_ ≈ 10^6^, reasonably consistent with the population size estimates from Ref.^32^

In summary, we have developed an ABC inference framework for simultaneous genome-wide prediction of selection strengths, mutation rates, and the fraction of viable alleles. The framework is based on the Ewens sampling formula, which we previously generalized to evolution under selection.^23^ Applying this approach to the evolutionary dynamics of a single *Drosophila* population, we observe, in line with previous reports, that a major fraction of the fly genome evolves under purifying selection against the constant influx of deleterious mutations. Moreover, we have found that genomic sequences can be classified into three distinct classes on the basis of their selection strength and investigated the effect of selection on mutation rate estimates. The accuracy of our predictions has been verified against synthetic data, which allowed us to systematically test all the major assumptions inherent in the model and gauge their potential effect on the accuracy of genome-wide predictions. Our computational approach can be used in other organisms for which population-level genomic data is available, providing an alternative to the Poisson Random Field and neutral approaches for explicit inference of key population-genetic parameters.

## Acknowledgements

We gratefully acknowledge stimulating discussions with Andrew Kern. AVM acknowledges support from the National Science Foundation (award MCB1920914).

## Supplementary Material

### Supplementary Figures

**Figure S1:**
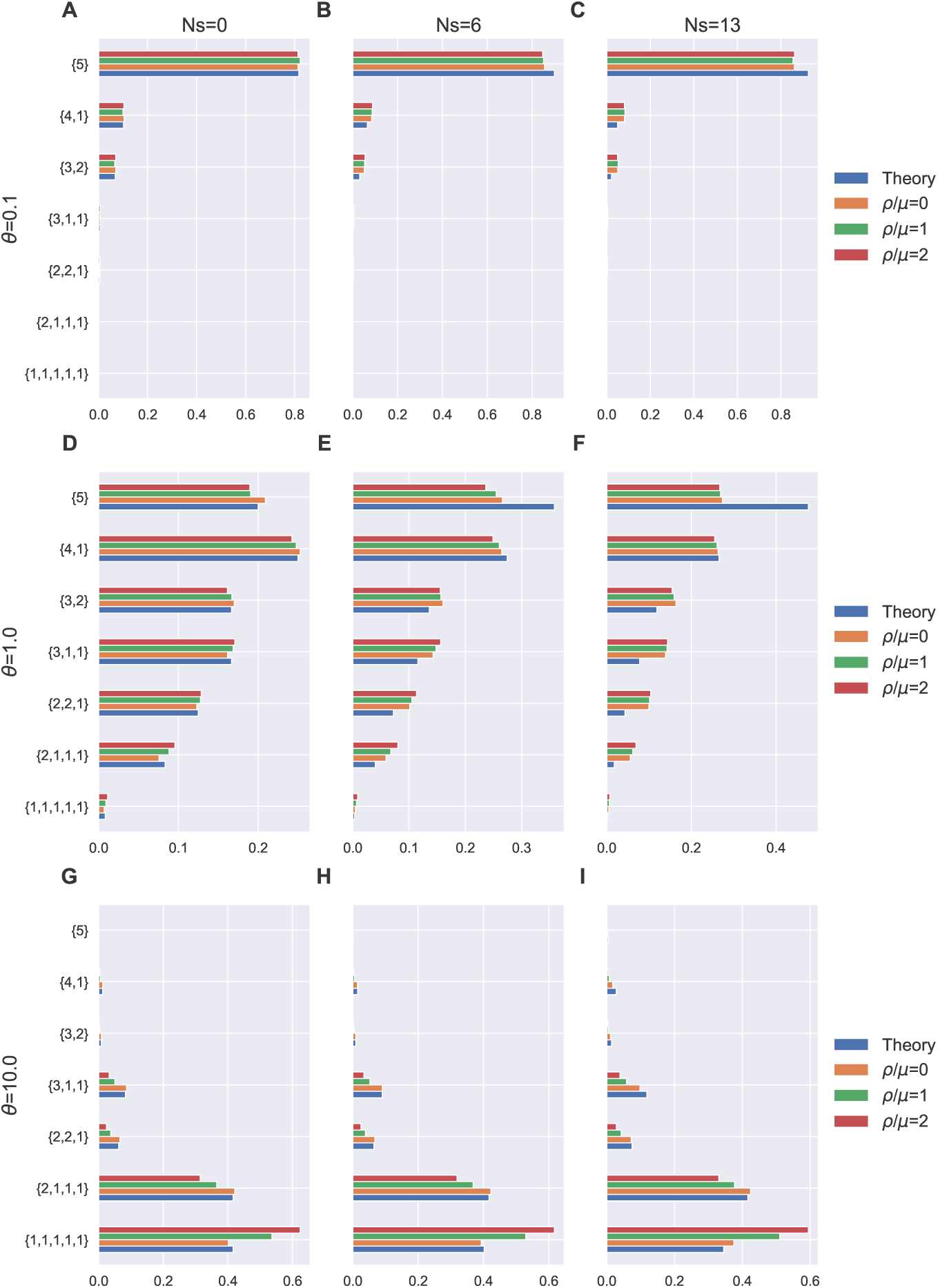
Effect of recombination on sampling frequencies. Columns of panels correspond to different selection strengths *Ns* and rows of panels to different values of the rescaled mutation rate *θ*. For cases with selection (*Ns* = 6 and 13), *γ* = 0.242 was used. In each panel, shown are sampling frequencies for the sample size *n′* = 5. Blue bars correspond to theoretical predictions (Eq. (1)), and orange, green and red bars correspond to numerical simulations averaged over 10^4^ independent steady-state samples (see Materials and Methods for details). Simulations with *ρ/µ* = 0: orange, *ρ/µ* = 1: green, *ρ/µ* = 2: red (*ρ* is the recombination rate and *µ* is the mutation rate per *L* = 10 locus).

**Figure S2:**
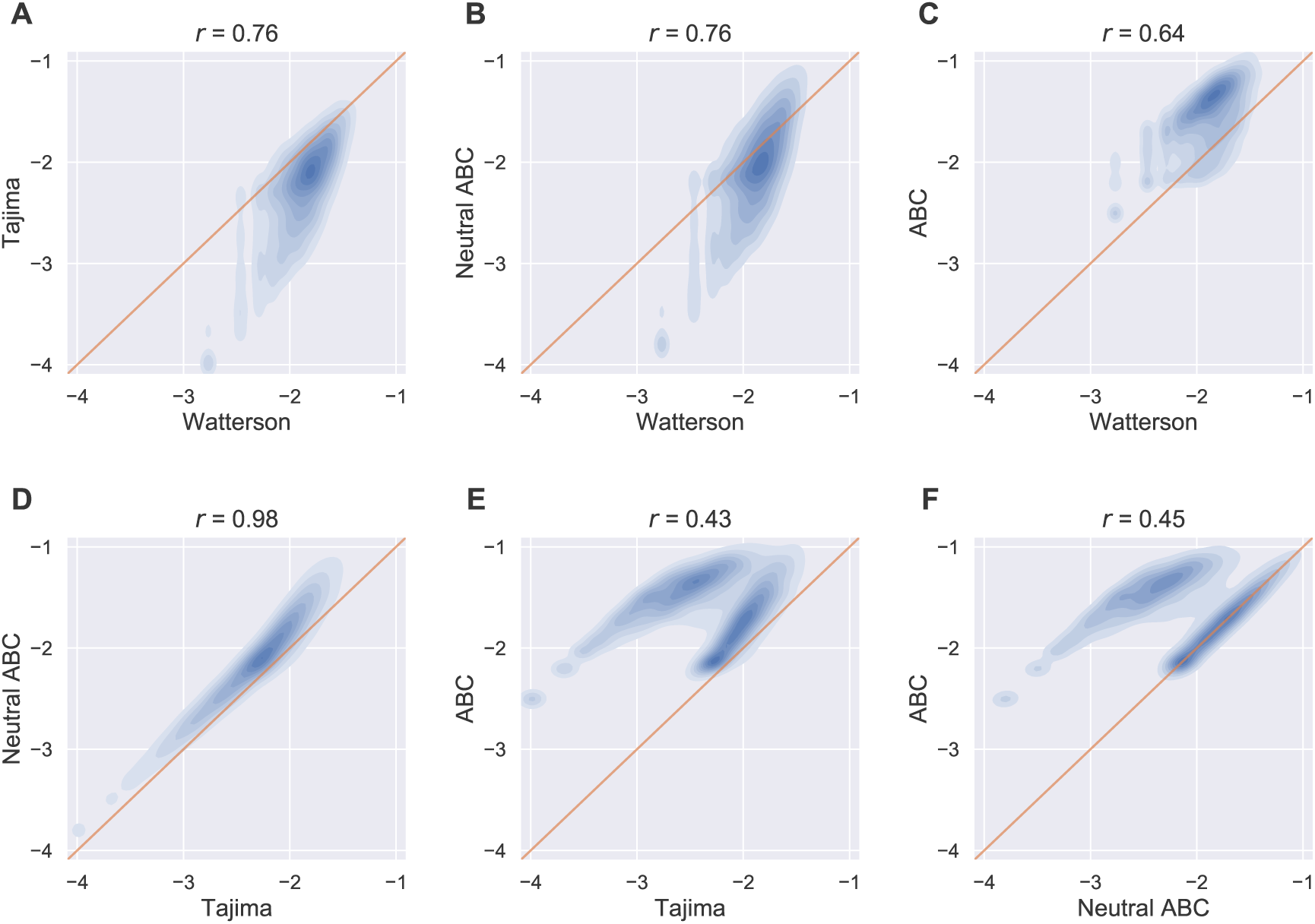
Comparison of mutation rate estimators. Shown are genome-wide density correlation plots (based on a random sample of 10^4^ windows) between Watterson and Tajima estimators (A), Watterson and neutral ABC estimators (B), Watterson and ABC with selection estimators (C), Tajima and neutral ABC estimators (D), Tajima and ABC with selection estimators (E), neutral ABC and ABC with selection (F). *r* is the linear correlation coefficient, computed for all windows. Both ABC estimators are represented by their median values in each genomic window. All estimators are shown on the log_10_ scale. Thin red lines in all panels have unit slopes.

**Figure S3:**
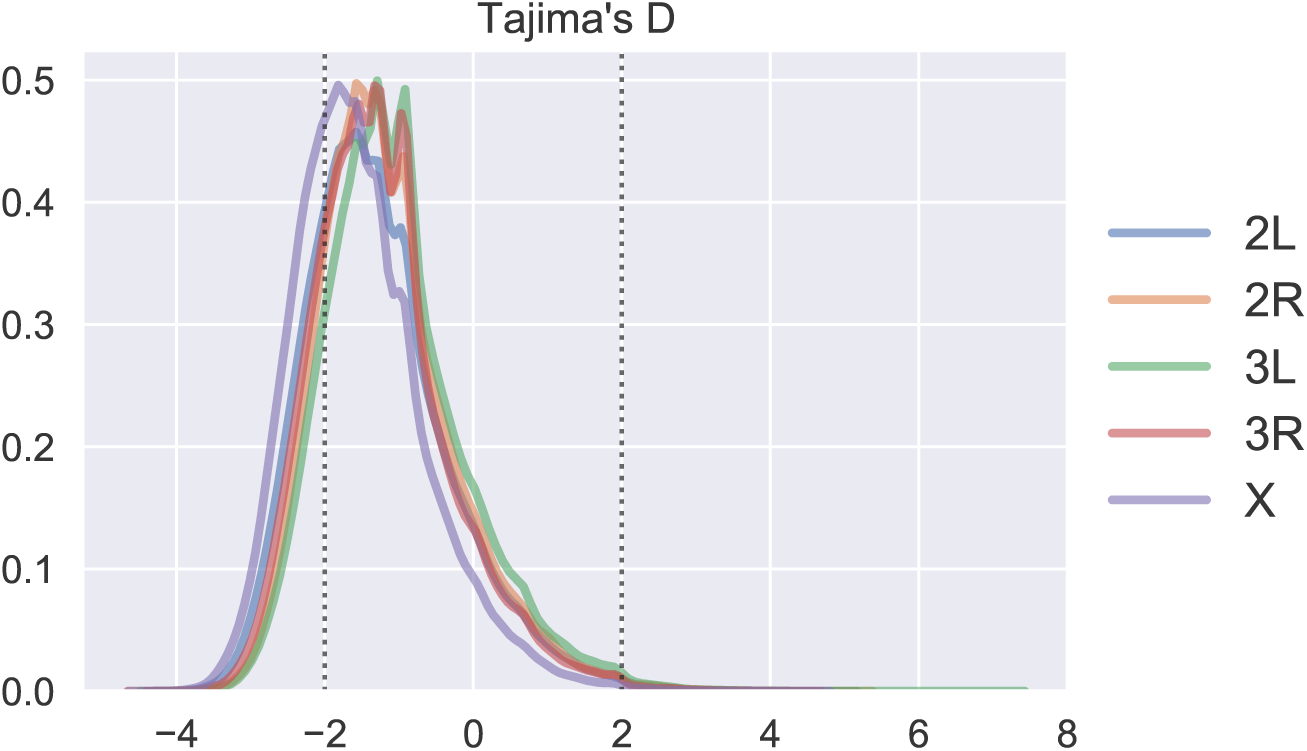
Chromosome-wide distributions of Tajima’s D statistic. Vertical dotted lines indicate *D* = −2 and *D* = 2; all windows with *D <* −2 or *D* > 2 are assumed to be under significant selection. The fractions of windows with *D <* −2 are 0.24, 0.17, 0.15, 0.20 and 0.32 for chromosomes 2L, 2R, 3L, 3R and X, respectively. The corresponding fractions of windows with *D* > 2 are 0.004, 0.004, 0.006, 0.003, 0.002.

**Figure S4:**
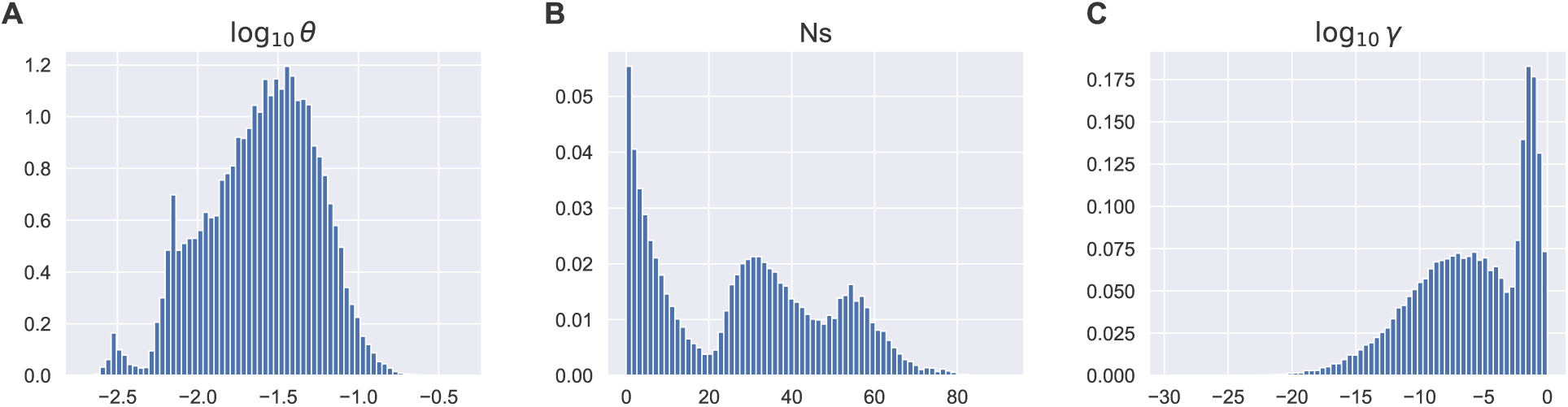
Genome-wide posterior distributions of mutation rates, selection coefficients, and fraction of viable genotypes. Each genome-wide posterior was created by combining individual posterior probabilities for each 100 bp window, which amounts to marginalizing all posteriors over the window index. (A): log_10_ *θ* distribution (*θ* is the rescaled mutation rate per bp), (B): *Ns* distribution, (C): log_10_ *γ* distribution.

**Figure S5:**
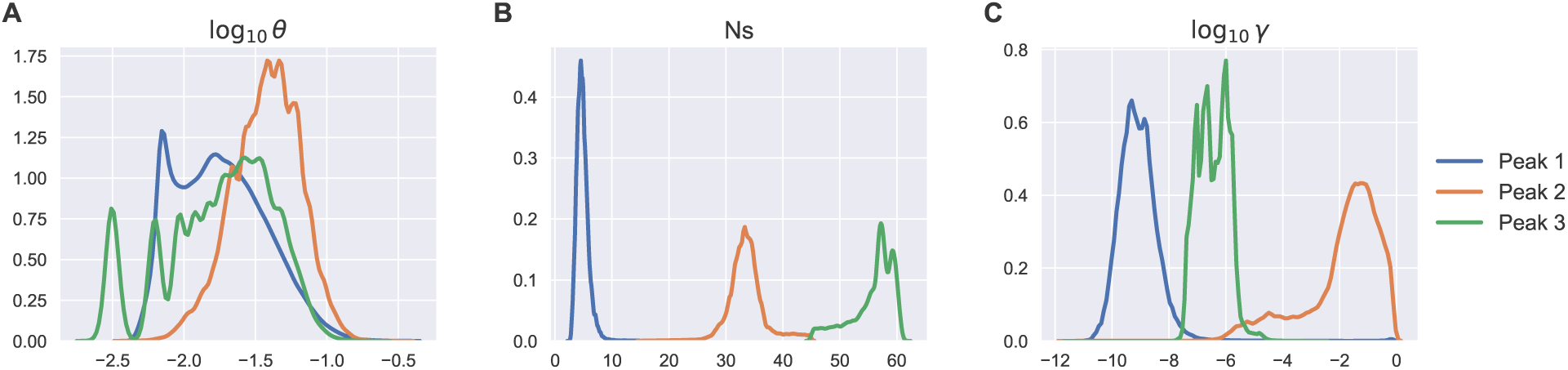
ABC inference of mutation rates, selection coefficients, and fraction of viable genotypes partitioned by selection strength. Genome-wide distributions of ABC inference results (median parameter values in each genomic window) for log_10_ *θ* (*θ* is the rescaled mutation rate per bp) (A), *Ns* (B) and log_10_ *γ* (C) partitioned by the range of *Ns*: *Ns <* 15 (blue), 15 ≤ *Ns* ≤ 45 (orange) and *Ns* > 45 (green). Each blue, orange and green curve is a smoothed trace of a normalized histogram.

**Figure S6:**
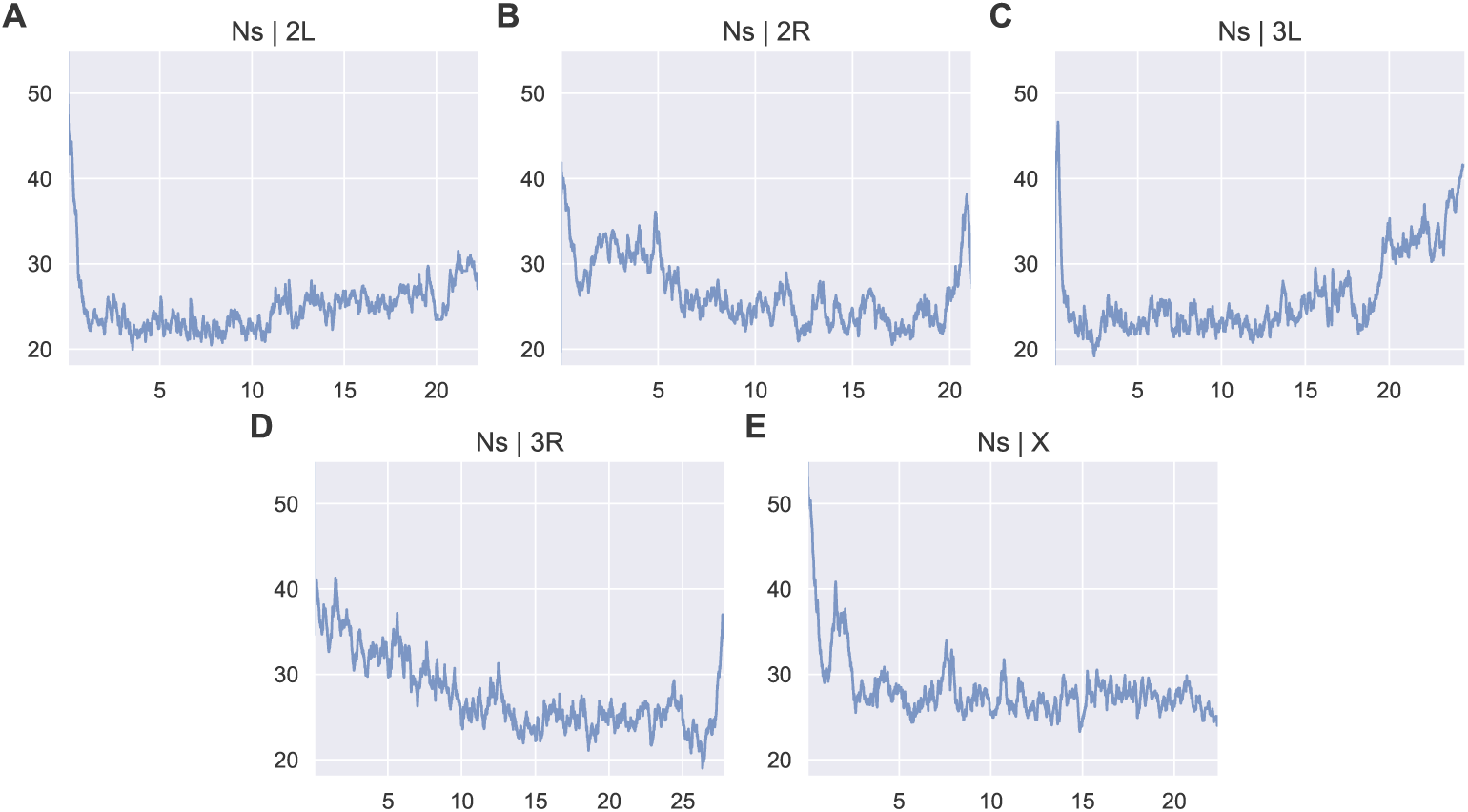
Inferred selection coefficients along *D. melanogaster* chromosomes. Shown are median values of selection coefficients (rescaled by the population size) predicted using ABC with selection, for each 100-bp window, for chromosomes 2L (A), 2R (B), 3L (C), 3R (D), and X (E), plotted vs. the genomic coordinate along each chromosome (in Mbp). All plotted values were smoothed with an exponentially weighted moving average with the center of mass of 1,000 windows, such that the exponential parameter *α* ≃ 10^−3^.

**Figure S7:**
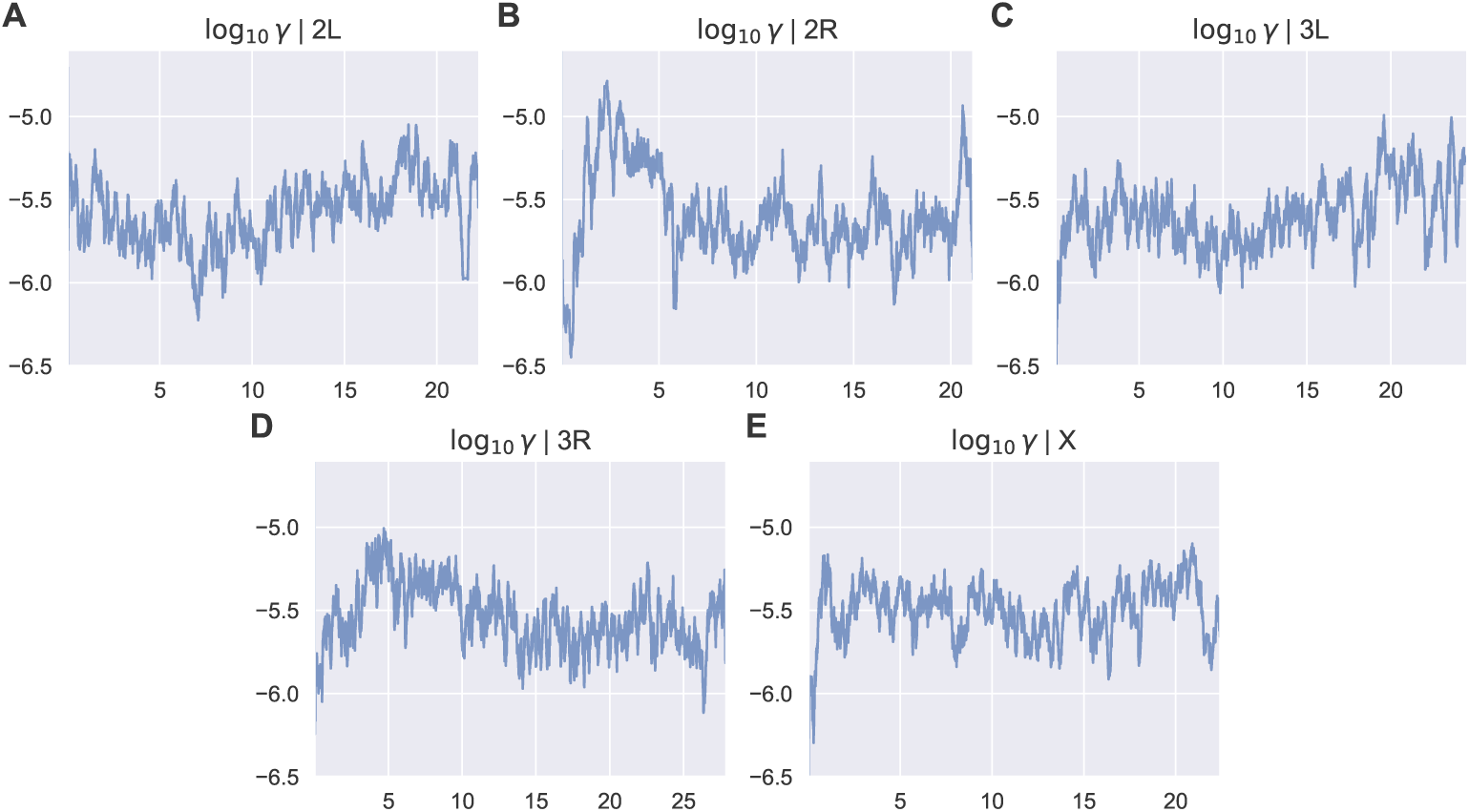
Inferred fractions of high-fitness alleles along *D. melanogaster* chromosomes. Shown are median values of the log fraction of viable genotypes, log_10_ *γ*, predicted using ABC with selection, for each 100-bp window, for chromosomes 2L (A), 2R (B), 3L (C), 3R (D), and X (E), plotted vs. the genomic coordinate along each chromosome (in Mbp). All plotted values were smoothed with an exponentially weighted moving average with the center of mass of 1,000 windows, such that the exponential parameter *α* ≃ 10^−3^.

**Figure S8:**
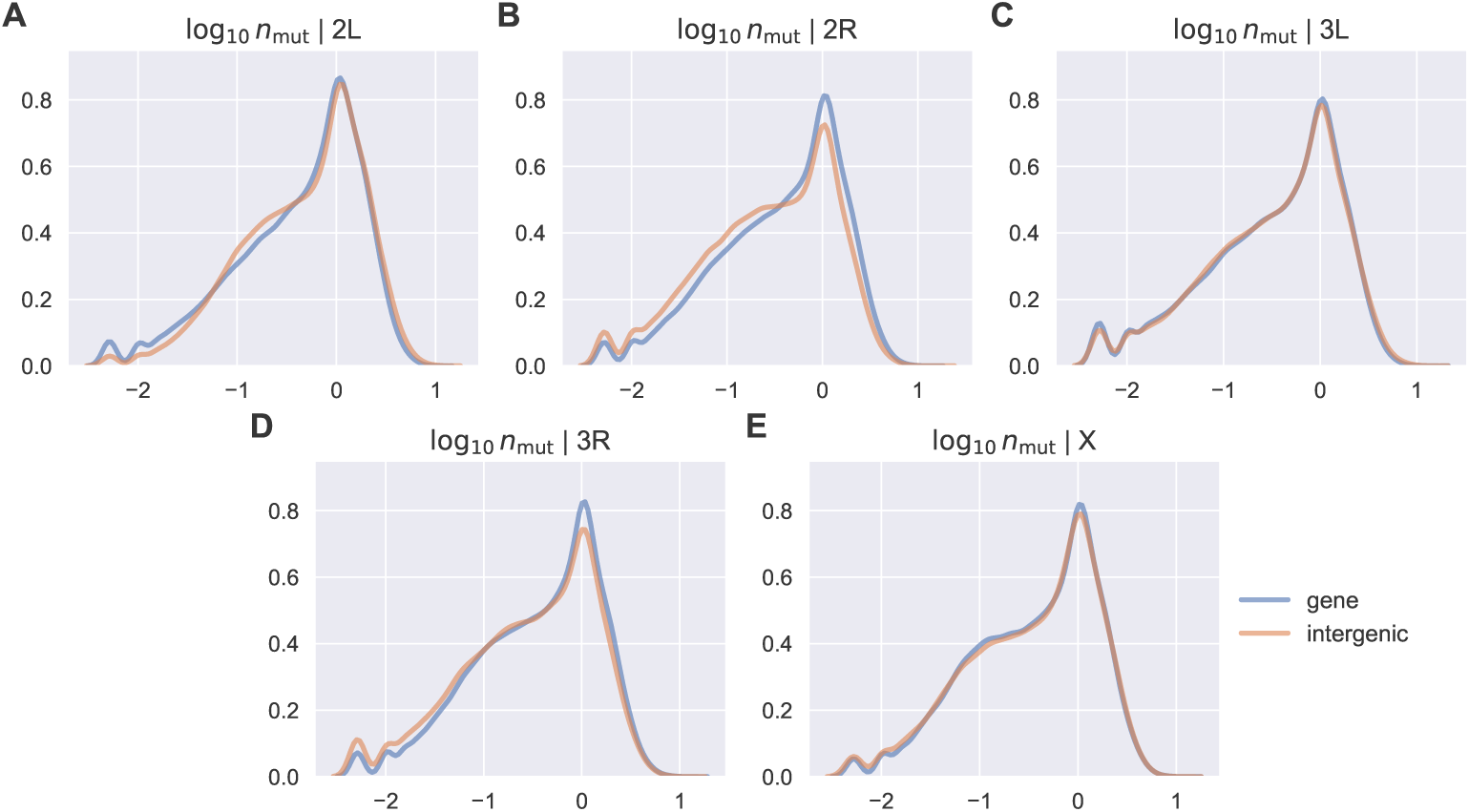
Chromosome-wide distributions of the number of mutations partitioned into functional regions. For each chromosome, shown is the average number of mutations per sequence in each alignment, *n*_mut_, for genes (blue) and intergenic regions (orange). Only windows that fully overlap the functional region of interest (gene or intergenic) are included. Panels A through E show the *n*_mut_ distributions for chromosomes 2L, 2R, 3L, 3R and X, respectively.

### Supplementary Tables

**Table S1:** ABC prediction accuracy in the presence of recombination. Shown are median values of the model parameters: *θ*_*m*_, (*Ns*)_*m*_, and *γ*_*m*_ predicted using the ABC inference pipeline. Sampling frequencies (*n′* = 5) for each set of parameters in Fig. S1 served as input to the algorithm. To facilitate comparisons, each subsection of the Table is marked by the corresponding Fig. S1 panel label.

| $\theta = 0.1, Ns = 0, \gamma = 0$ | | | | |
| --- | --- | --- | --- | --- |
| **A** | Theory | $\rho/\mu = 0$ | $\rho/\mu = 1$ | $\rho/\mu = 2$ |
| $\theta_m$ | 0.10 | 0.10 | 0.10 | 0.10 |
| $(Ns)_m$ | 4.59 | 4.21 | 4.62 | 4.01 |
| $\gamma_m$ | $1.24 \times 10^{-7}$ | $1.55 \times 10^{-9}$ | $2.00 \times 10^{-6}$ | $2.67 \times 10^{-9}$ |
| $\theta = 0.1, Ns = 6, \gamma = 0.24$ | | | | |
| **B** | Theory | $\rho/\mu = 0$ | $\rho/\mu = 1$ | $\rho/\mu = 2$ |
| $\theta_m$ | 0.12 | 0.10 | 0.10 | 0.10 |
| $(Ns)_m$ | 11.92 | 9.08 | 5.08 | 6.56 |
| $\gamma_m$ | 0.23 | 0.58 | 0.02 | 0.53 |
| $\theta = 0.1, Ns = 13, \gamma = 0.24$ | | | | |
| **C** | Theory | $\rho/\mu = 0$ | $\rho/\mu = 1$ | $\rho/\mu = 2$ |
| $\theta_m$ | 0.11 | 0.09 | 0.09 | 0.10 |
| $(Ns)_m$ | 13.85 | 12.95 | 5.62 | 16.18 |
| $\gamma_m$ | 0.20 | 0.59 | 0.16 | 0.61 |
| $\theta = 1.0, Ns = 0, \gamma = 0$ | | | | |
| **D** | Theory | $\rho/\mu = 0$ | $\rho/\mu = 1$ | $\rho/\mu = 2$ |
| $\theta_m$ | 1.00 | 0.96 | 1.03 | 1.10 |
| $(Ns)_m$ | 4.66 | 5.25 | 5.01 | 9.30 |
| $\gamma_m$ | $4.59 \times 10^{-8}$ | $2.79 \times 10^{-9}$ | $4.13 \times 10^{-9}$ | $2.30 \times 10^{-5}$ |
| $\theta = 1.0, Ns = 6, \gamma = 0.24$ | | | | |
| **E** | Theory | $\rho/\mu = 0$ | $\rho/\mu = 1$ | $\rho/\mu = 2$ |
| $\theta_m$ | 1.07 | 1.01 | 1.00 | 1.04 |
| $(Ns)_m$ | 7.19 | 8.55 | 3.81 | 9.46 |
| $\gamma_m$ | 0.27 | 0.58 | 0.09 | $8.80 \times 10^{-5}$ |
| $\theta = 1.0, Ns = 13, \gamma = 0.24$ | | | | |
| **F** | Theory | $\rho/\mu = 0$ | $\rho/\mu = 1$ | $\rho/\mu = 2$ |
| $\theta_m$ | 1.12 | 0.89 | 0.92 | 0.97 |
| $(Ns)_m$ | 16.29 | 5.66 | 4.44 | 7.91 |
| $\gamma_m$ | 0.22 | 0.54 | 0.03 | $4.48 \times 10^{-4}$ |

Table S1: **ABC prediction accuracy in the presence of recombination.**
| $\theta = 10.0, Ns = 0, \gamma = 0$ | | | | |
| --- | --- | --- | --- | --- |
| **G** | **Theory** | $\rho/\mu = 0$ | $\rho/\mu = 1$ | $\rho/\mu = 2$ |
| $\theta_m$ | 10.00 | 9.59 | 14.76 | 20.30 |
| $(Ns)_m$ | 7.11 | 5.76 | 10.72 | 14.40 |
| $\gamma_m$ | $1.00 \times 10^{-6}$ | $1.88 \times 10^{-7}$ | 0.03 | 0.06 |
| $\theta = 10.0, Ns = 6, \gamma = 0.24$ | | | | |
| **H** | **Theory** | $\rho/\mu = 0$ | $\rho/\mu = 1$ | $\rho/\mu = 2$ |
| $\theta_m$ | 9.86 | 9.31 | 14.56 | 19.61 |
| $(Ns)_m$ | 10.03 | 7.73 | 13.25 | 14.21 |
| $\gamma_m$ | 0.04 | $5.00 \times 10^{-6}$ | 0.01 | 0.02 |
| $\theta = 10.0, Ns = 13, \gamma = 0.24$ | | | | |
| **I** | **Theory** | $\rho/\mu = 0$ | $\rho/\mu = 1$ | $\rho/\mu = 2$ |
| $\theta_m$ | 9.89 | 8.91 | 13.83 | 18.48 |
| $(Ns)_m$ | 12.44 | 8.65 | 11.18 | 16.82 |
| $\gamma_m$ | 0.24 | 0.01 | 0.07 | 0.03 |

**Table S2:**
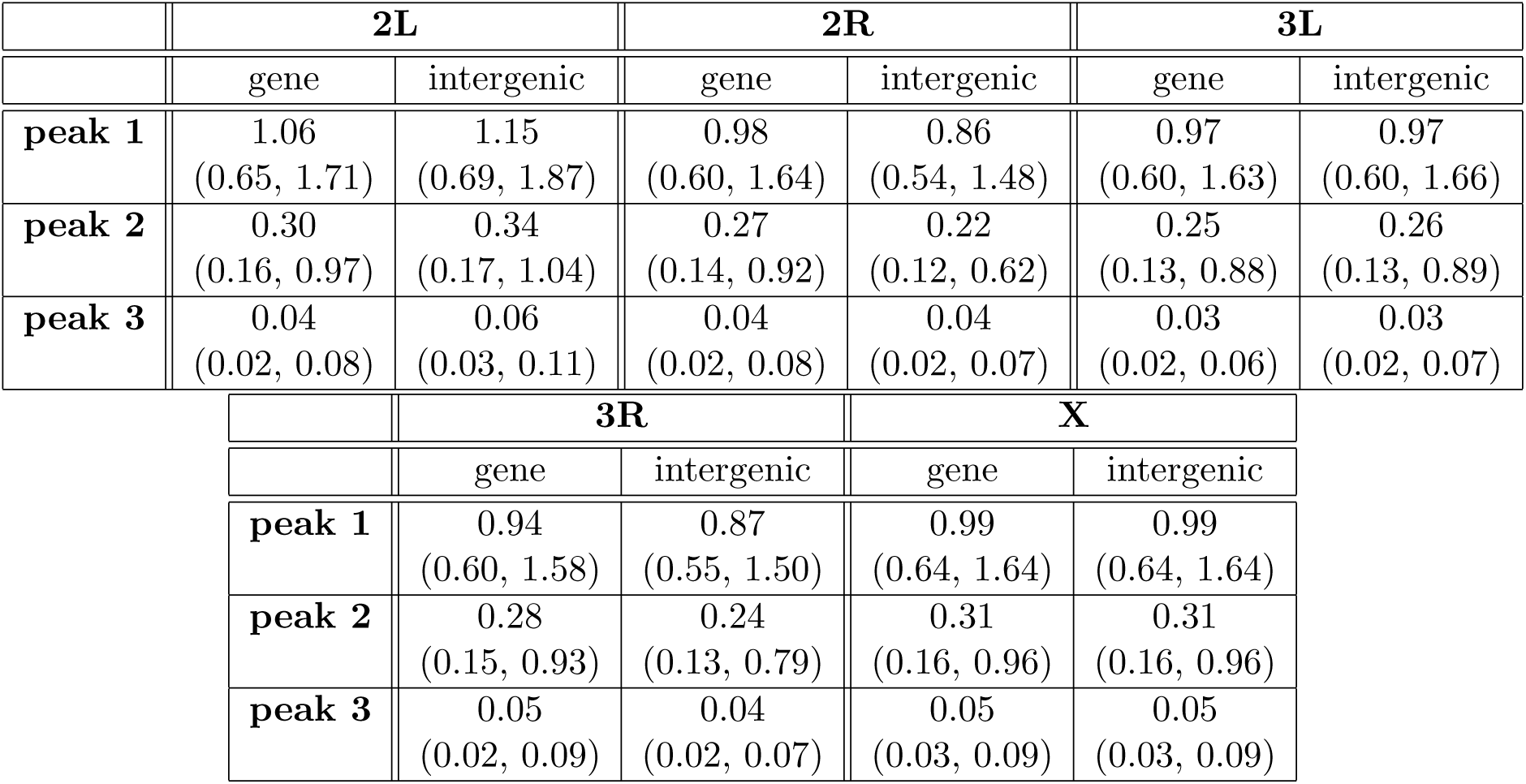
Summary statistics for the average number of mutations per sequence, partitioned by functional region and selection strength. Shown are the median values, followed by the first quartile and the third quartile in parentheses, of the distribution of *n*_mut_, the average number of mutations per sequence in each 100 bp window, for all windows in a given chromosome, sorted by the selection peaks in Fig. 5B and by the functional region (gene or intergenic).

